# Persistent Gamma Spiking in Non-Sensory Fast-Spiking Cells Predicts Perceptual Success

**DOI:** 10.1101/374314

**Authors:** Hyeyoung Shin, Christopher I. Moore

## Abstract

Persistent gamma oscillations (30-55Hz) are hypothesized to temporally coordinate stimulus encoding, enabling perception. This prediction poses a conundrum: How can gamma serve as a template when the stimulus itself drives its mediators, presumably perturbing its maintenance? Specifically, fast-spiking interneurons (FS), a key gamma generator, can be highly sensory responsive. Further, the gamma-band local field potential (LFP) shows properties inconsistent with temporal coordination. Combining tetrode recording with controlled psychophysics revealed an FS subtype (γnsFS) that was not sensory responsive, whose inter-spike intervals peaked at gamma, and that fired with higher periodicity than other FS. Successful detection was predicted by increased regularity in γnsFS spiking at gamma, persisting from before to after sensory onset. In contrast, gamma LFP power negatively predicted detection, and was negatively related to gamma band spiking by γnsFS. These results suggest that a distinct interneuron subgroup, not ‘distracted’ by sensory input, mediates perceptually-relevant oscillations independent of LFP.

## Introduction

Neocortical gamma oscillations are evident during behaviorally-relevant neural computations including perception (Fries et al., 2001; Gray et al., 1989; Siegle et al., 2014), cognition (Cho et al., 2015; Kim et al., 2016) and action (Niell and Stryker, 2010). Decreases in gamma expression are a biomarker for numerous neurological disorders (Iaccarino et al., 2016; Lewis et al., 2005, 2012; Uhlhaas et al., 2008). These studies suggest a role for gamma, or its component mechanistic processes, in successful information processing. However, there is intense debate whether this dynamic plays a meaningful computational role, or is simply a by-product of local resonant excitatory-inhibitory circuits that are active during such behaviors (Ray and Maunsell, 2010, 2015; Shadlen and Movshon, 1999; Xing et al., 2012).

Hypotheses proposing a role for gamma in sensory neocortical relay fall into two broad categories (Knoblich et al., 2010; Pritchett et al., 2015). First, gamma could enhance relay by improving rate coding locally; for example by increasing firing in appropriate projection neurons (Azouz and Gray, 2000; Fellous et al., 2003; Tiesinga et al., 2004); or through effective inhibition of distracting ‘surround’ neurons that carry non-informative signals (Gieselmann and Thiele, 2008; Vinck and Bosman, 2016); or both (Borgers et al., 2008). In these scenarios, the exact frequency and duration of expression could vary within and between areas while still achieving effective rate enhancement (Pritchett et al., 2015).

Much of the interest in gamma stems from a second class of hypotheses that predict that this rhythmic process coordinates action potential timing across a long-range network (Gray et al., 1989; Singer, 1993; Singer and Gray, 1995). The coordination of timing across brain areas has been hypothesized to ‘bind’ the activity of participating cells into a coherent perception of a single entity. Aligned timing could in turn enhance the efficacy of disparate elements in driving a common target. A related proposal, ‘communication through coherence,’ has suggested that relay between areas relies on phase alignment of gamma in sending and receiving representations (Fries, 2009, 2015). Central to binding hypotheses is the existence of a ‘reference’ gamma that emerges prior to, and persists unperturbed by, stimulus presentation. This persistent reference gamma can then organize distinct, non-local representations, independent of specific stimulus features (Singer, 1993; Singer and Gray, 1995).

Several challenges have been raised to the binding-by-gamma view. Using the local field potential (LFP) as a gamma metric, studies in visual neocortex have shown that subtle changes in features that are not relevant for identifying stimulus identity, such as contrast, can change gamma frequency and power (Ray and Maunsell, 2010). This sensitivity of the LFP gamma to subtle changes in specific stimulus features undermines the possibility that a sustained coordinating reference can be indexed by the LFP gamma. Moreover, differences in LFP frequency and power can be observed across positions in a visual map driven by a common object, further undermining the likelihood of lateral binding (Ray and Maunsell, 2010, 2015).

Another logical challenge to the binding hypothesis concerns the mechanisms of gamma origin. Neocortical gamma is typically found to depend on coordinated firing of fast-spiking interneurons (FS) that create highly effective inhibition lasting ~20 ms (Buzsáki and Wang, 2012; Tiesinga and Sejnowski, 2009; Whittington et al., 2000; but see also Veit et al., 2017). This period effectively sets the duration of a gamma cycle, creating a brief window of opportunity for more synchronous firing of projection neurons upon relaxation from hyperpolarization. Such synchronization of projection neurons can lead to an efficient relay of signals to downstream targets, and the onset of another gamma cycle (Borgers and Kopell, 2003; Cardin et al., 2009; Chen et al., 2017; Moore et al., 2010; Sohal et al., 2009). Within neocortical areas such as primary somatosensory ‘barrel’ neocortex (SI), FS are typically found to be the most sensitive neurons to sensory stimulation, responding at short latencies and with high consistency to weak peripheral inputs (Andermann et al., 2004; Cruikshank et al., 2007; Simons and Carvell, 1989; Swadlow, 2003). Some studies have reported that all FS encountered in SI are sensory responsive (Simons, 1978; Swadlow, 2003). This sensitivity to sensory drive suggests that FS could not sustain a reference gamma rhythm initially established prestimulus, as local FS firing patterns should be disrupted by the input they seek to temporally organize, altering the reference gamma rhythm. In sum, inconsistencies between LFP recording and binding hypothesis predictions, and the high degree of FS sensitivity in sensory neocortex, argue against a role for gamma in temporal coordination across sensory neocortices.

To understand the endogenous behavior of FS during sensory processing, we recorded extensively from FS in SI using chronic tetrode implants while mice performed well-controlled vibrissal motion detection, a task previously shown to benefit from the optogenetic induction of FS synchrony at gamma (Siegle et al., 2014). We found that the majority of FS were non-sensory, failing to show a significant change in rate in response to sensory stimulation. These non-sensory FS differentiated detected (hit) trials versus non-detected (miss) trials with greater inter-spike interval (ISI) regularity in the gamma range. The rhythmic spiking of these γ-spiking non-sensory FS neurons (γnsFS) persisted from prestimulus to poststimulus without interruption, and was not reset by sensory drive. While γnsFS spiking at gamma positively predicted detection, their firing was negatively correlated with gamma band LFP power, which in turn negatively predicted detection success. In contrast, gamma band LFP power was positively correlated with the spike gamma power of sensory-responsive FS (sFS), suggesting that this FS subtype regulates large pyramidal neurons en masse. Poststimulus LFP and sFS gamma power linearly correlated with the stimulus amplitude, whereas spike gamma power of γnsFS and plnsFS had no relationship with the stimulus amplitude. In sum, these findings show the existence of a distinct FS subtype, γnsFS, that preferentially demonstrate gamma period ISIs, and this gamma spiking pattern is further enhanced during perceptually successful trials. This FS subtype could provide a reference signal for temporal coordination, independent of the LFP and specific stimulus features.

## Results

To investigate the relationship between endogenous FS activity and perception, we recorded mice trained to respond to vibrissal deflection using chronic tetrode implants in SI (see *Methods*). Data presented are from sessions with high-quality psychometric behavior (4 mice, 128 sessions; **Supplemental Figure 1A** and *Methods*). Water was delivered if detection was reported ≤700 ms after sensory onset. Catch trials were trials where the stimulus amplitude was zero. Catch trials were implemented to approximate what percentage of hit trials the animal ‘guessed’ correctly by chance (Macmillan and Creelman, 2004). Incorrect responses during a catch window are referred to as false alarms, and led to a 15 s timeout from the task. We compared detected (hit) and non-detected (miss) trials matched in stimulus amplitude (**Supplemental Figure 1B**; see *Methods*). In analyzing matched hit trials, we limited selection to those with reaction times >100 ms, reflecting the point at which there was a substantial initial increase in reaction times (**Supplemental Figure 1C**). Spike width, indexed as the time between peak and trough of an action potential, provided high separability of FS (N=188) from the broader population of putative excitatory neurons (**Supplemental Figure 1D**). We further grouped FS into sensory responsive (sFS) and non-sensory responsive (nsFS), based on an ideal observer analysis for discrimination of 0 – 100 ms firing rate on maximal stimulus amplitude trials versus zero amplitude catch trials (**Supplemental Figure 1E** and *Methods*).

### A Minority of FS are Sensory Responsive (sFS), and these sFS Predict Successful Detection with Lower Rates Prestimulus and Higher Rates Poststimulus

Consistent with the sparse probability that SI neurons in general respond to controlled vibrissal deflections (Kwon et al., 2016; Simons and Carvell, 1989), sFS constituted 36.7 % of the recorded FS population (N=69 sFS of 188 FS). Among sFS, differences in rate predicted successful detection in distinct ways pre- and poststimulus (**Supplemental Figure 1F**). Prestimulus, the sFS subtype showed lower activity on hits compared to misses (spike rate from −1000 to 0 ms relative to sensory onset, mean ± SEM: Hits 10.82 ± 1.00 Hz; Misses 12.23 ± 1.04 Hz; p=9.80×10^−6^ Wilcoxon signed rank test). This difference inverted at sensory onset, with stronger sensory evoked responses on hits. For all time points >5 ms poststimulus, sFS rates were higher on hits than on misses (spike rate 0 – 100 ms relative to sensory onset, mean ± SEM: Hits 18.39 ± 1.40 Hz; Misses 15.36± 1.14 Hz; p=1.34×10^−7^). Evoked rate differences between hits versus miss trials were significant across the entire psychometric range (**Supplemental Figure 1G**).

### A Distinct Subtype of Non-Sensory FS (γnsFS) Spike Most Frequently at Gamma Intervals (~25 ms)

To begin to investigate the relationship between the pattern of FS spiking and perceptual performance, we calculated FS inter-spike intervals (ISIs) during the baseline period. **Figure 1A** shows the distribution of the most common ISIs for each FS (i.e., the peak of the ISI distribution calculated in 5 ms bins, 1 ms steps). Most sFS (***gray bars***) showed peak ISIs ≤18 ms (N=53/69 neurons). In contrast, non-sensory responsive FS (nsFS; ***colored bars***) showed a segregation of peak ISIs into two groups, defined by a gap in the distributions from 15 to 18 ms. One group showed peak ISIs ≤14 ms (N=54), and the other peak ISIs >18 ms (N=60), and primarily between 18 – 33 ms. This range of peak ISIs corresponds to a period in a ~30 – 55 Hz gamma oscillation, also known as the ‘low’ gamma-band.

The short-duration ISI peaks in sFS, and in a subtype of nsFS, are consistent with a Poisson-like process with a spike refractory period (~5 ms in most neurons; Kass et al., 2014). **Figure 1B** shows the ISI distributions for the three subtypes defined by their peak ISIs. The sFS distribution showed Poisson-like firing behavior (***gray***), as did the nsFS subtype with peak ISIs <14 ms (***olive***). For ease of reference, we will term this nsFS subtype Poisson-like (**plnsFS**). The other nsFS subtype will be referred to as the gamma-nsFS (**γnsFS**), as they showed the highest rate of peak ISIs at ~25 ms, corresponding to a ~40 Hz frequency (***teal***).

### γnsFS Spike More Periodically than Other Subtypes

The nsFS ISIs distributions indicate two distinct subtypes, with one (γnsFS) showing a propensity to fire in the gamma range. However, the likelihood of demonstrating this ISI range does not indicate the extent to which the spiking is periodic. To examine spiking regularity, we measured the coefficient of variation (CV^2^; the ISI variance divided by the mean squared ISI). For ISIs sampled over a sufficiently long time window, a Poisson process has a CV^2^ of 1, with higher values indicating more variable ISIs (e.g., less rhythmic spiking). **Figure 1C** shows the histogram of CV^2^ across all three FS subtypes. Most plnsFS showed CV^2^>1 (N=48/54; mean ± SEM: 1.221 ± 0.036) as did most sFS (N=62/69; mean ± SEM: 1.375 ± 0.0776). In contrast, most γnsFS showed CV^2^<1 (N=51/60; mean ± SEM: 0.937 ± 0.012). These data indicate that γnsFS are not only separable from other subtypes by spiking more often at gamma intervals, but also by spiking with greater regularity. Separation of this subtype can be appreciated in **Figure 1D**, which shows the relationship between CV^2^ and ISI in the prestimulus period for each of the three subtypes. Figure 1E shows autocorrelograms from example units of each FS subtype.

These differences in baseline spike properties between γnsFS and sFS suggests that these groups have distinct biophysical properties and/or afferent connectivity, and that these differences may contribute to their sensory responsiveness. Unlike γnsFS, plnsFS spiking properties were similar to sFS. This similarity suggests that plnsFS may be non-sensory in our experimental conditions because of our use of a sensory stimulus that varied on a single direction axis, frequency and duration.

In the psychophysical paradigm employed here, we chose to stimulate all macrovibrissae posterior to the 4 arc, to minimize the chance of misalignment between our tetrode implants and recording sites. Nevertheless, regional differences within the SI macrovibrissal representation can change the probability of sensory response (e.g., septal alignment, differences in depth, direction preference columns), and nsFS would be predicted to occur more commonly in positions that were generally less responsive. To test for the possibility that the differences in sFS and the nsFS subtypes resulted solely from tetrode positioning, we limited analyses to FS recorded from tetrodes where robust sensory evoked multi-unit activity (MUA) was observed. All differences between sFS and the nsFS subtypes described above were observed when analyses were limited to these significantly sensory responsive tetrodes (**Supplemental Figure 2A**). In addition, we also tested the possibility that the difference between sFS and nsFS subtypes depended on the specific definition of sensory responsiveness that we employed. We compared two additional definitions and found the effects described above to be robust across these definitions (**Supplemental Figures 2B-C**).

### Regular Spiking at Gamma by γnsFS Before Sensory Onset Predicts Successful Performance

We next tested whether spiking patterns in the three FS subtypes predicted sensory performance. We first compared the incidence of gamma range ISIs in the prestimulus period on hit versus miss trials. As shown in **Figure 1F *left***, the sFS subtype showed a lower occurrence gamma ISIs on hit versus miss trials (count of 18 – 33 ms ISIs on hit versus miss trials; p=4.32×10^−5^ Wilcoxon signed rank test), consistent with their lower prestimulus firing rate on hits (**Supplemental Figure 1F**). In contrast, γnsFS showed an increase in the likelihood of gamma range ISIs on hit versus miss trials (**Figure 1F *center*** p=0.0145), accompanied by higher prestimulus firing rate on hits (**Supplemental Figure 1F**).

All three subtypes showed greater regularity (lower CV^2^) in the prestimulus period on hits versus miss trials (**Figure 1Gi**; sFS p=5.51χ10^−5^; γnsFS p=6.40χ10^−6^; plnsFS p=2.17×10^−4^ Wilcoxon signed rank test). However, this lower CV^2^ emerged for different reasons in the different subtypes. For sFS, the mean ISI increased on hits (**Figure 1Gii**; p=9.89χ10^−7^), leading to lower CV^2^ despite also showing larger variance on hits than misses (**Figure 1 Giii**; p=0.015). These mean ISI and rate differences between hits and misses in sFS were paralleled by similar trends in the plnsFS (**Figure 1Gii**, mean ISI difference p=0.089; **Supplemental Figure 1F**, prestimulus rate difference p=0.030). ISI variance was not different between hits and misses for plnsFS (**Figure 1Giii**; p=0.198). In contrast, γnsFS showed no change in mean ISI (**Figure 1Gii**; p=0.503), reflecting its relative constancy in gamma interval spiking. Instead, γnsFS showed a decrease in variance on hits relative to misses (**Figure 1Giii**; p=0.044), consistent with more regular spiking patterns on hit trials.

As shown in **Figure 1B**, plnsFS typically showed shorter ISIs than γnsFS. However, on hit trials, their rate decreased, which was associated with the emergence of a concentration of cells with peak ISIs in the gamma range (**Figure 1H**). For the plnsFS, this shift in peak location was significant (p=1.88χ10^−6^ Wilcoxon signed rank test). Peak ISIs did not shift detectably among γnsFS or sFS (sFS p=0.687; γnsFS p=0.201).

In sum, these data show that successful sensory performance is predicted by a prestimulus increase in the regularity of γnsFS spiking, and their increase in gamma ISIs. In contrast, sFS showed higher variance on hits and a lower occurrence of ISIs over a broad range including gamma.

### Enhanced Gamma Interval Spiking by γnsFS Persists from Pre-to Poststimulus

We next asked whether the predictive gamma spiking behavior of γnsFS persisted into the poststimulus period. To test this possibility, we plotted ISI density heat maps for the period surrounding sensory stimulation on hits, misses and their difference (**Figure 2A**). For sFS and plnsFS, the Poisson-like fall-off with increasing ISIs in the prestimulus period (as shown in **Figure 1F**) is evident in these plots as a shift from red to blue from the bottom (short ISIs) to the top (longer ISIs) of each plot. In the hit minus miss map for sFS, lower density of ISIs on detected trials is evident in the prestimulus period (*blue* prior to stimulus onset), reflecting a lower overall rate (**Supplemental Figure 1F**). Poststimulus, a sharp increase in ISI densities was observed *(red* immediately poststimulus), most prominently for short ISIs, again corresponding to the rate differences described above (**Supplemental Figure 1F**).

**Figure 1.**
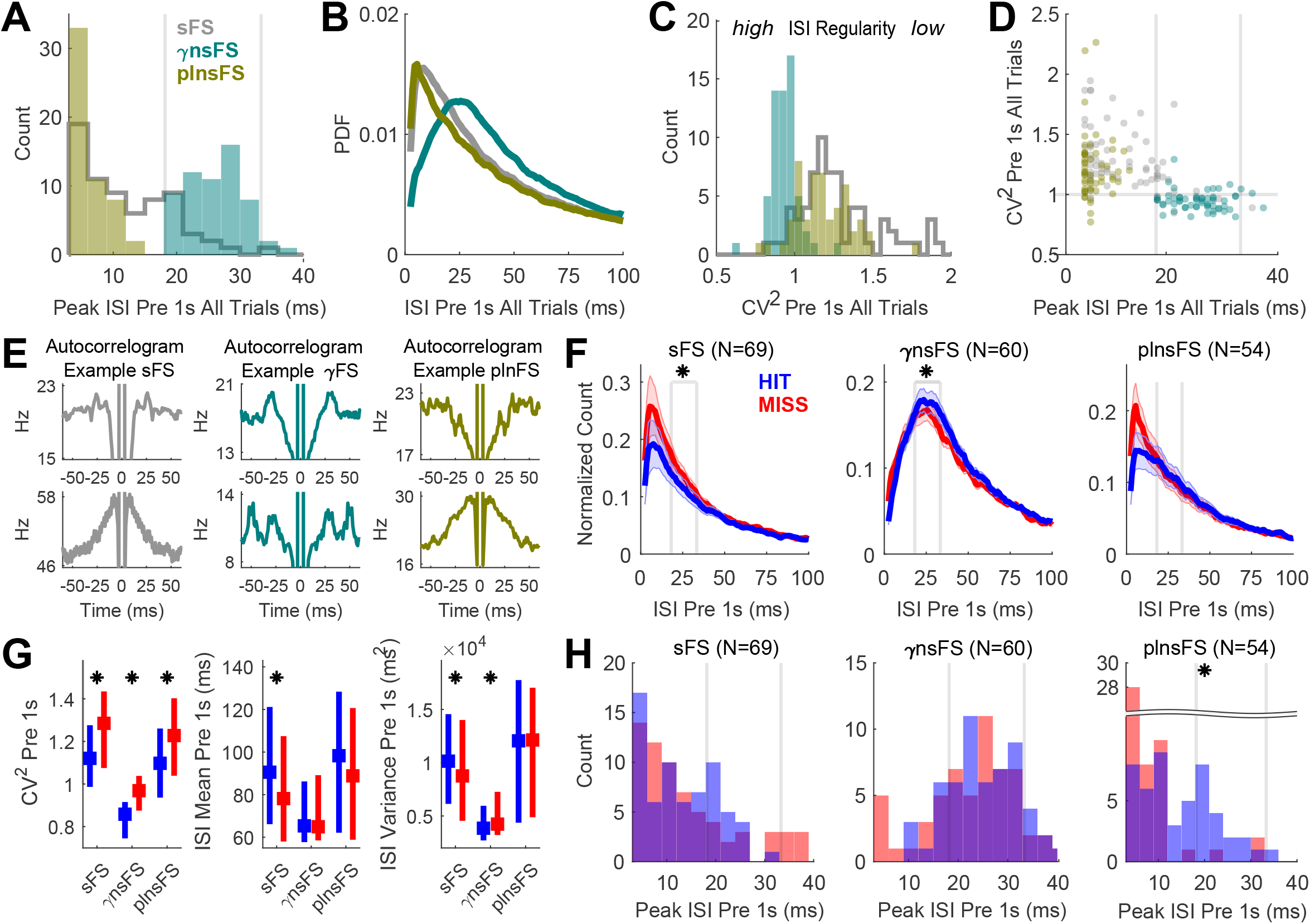
A Distinct Subset of Non-Sensory Responsive FS Spiked Most Frequently at Gamma Intervals, was More Periodic than Other FS, and Predicted Successful Performance by Enhanced Gamma Interval Spiking. **A-D.** sFS *gray*; γnsFS *teal*; plnsFS *olive*. Inter-spike interval (ISI) distribution and coefficient of variance squared (CV^2^) were calculated for each FS, from ISIs in the 1 s prestimulus period, pooled across all trials. ISI probability distribution function (PDF) was calculated in 5 ms bins, sliding in 1 ms steps. **A.** The peak of inter-spike interval (ISI) distributions was calculated for each FS. The histogram of ISI peaks (3 ms bins) for non-sensory responsive FS (nsFS) reveal a gap between 15 – 18 ms, separating their firing patterns into two subtypes. The subtype with peak ISIs predominantly between 18 – 33 ms was named ‘gamma spiking’ non-sensory FS (γnsFS; *teal*), reflecting the overlap in spike intervals and gamma frequencies (~30 – 55 Hz). Those with peaks at faster ISI intervals (<15 ms) showed Poisson-like firing patterns (plnsFS; *olive*). The sensory responsive FS (sFS; *gray*) also showed Poisson-like firing patterns. **B.** The ISI probability distribution function (PDF) was averaged across FS within each subtype, revealing Poisson-like distributions for plnsFS and sFS, and a gamma-band centered distribution for γnsFS. **C.** The histogram of CV^2^ values (bin size 0.05) showed greater regularity in baseline firing for γnsFS, compared to plnsFS or sFS. **D.** FS subtypes were separable in the scatterplot of ISI peak and CV^2^ for all FS. Most γnsFS had CV^2^<1 and peak ISIs in the gamma range, whereas most plnsFS and sFS had CV^2^>1 and peak ISIs<15 ms. **E.** Example autocorrelograms from each FS subtype (1 ms bins, smoothed with 5 ms boxcar window). **F-H.** Hit trials *blue;* miss trials *red*. **F.** Histograms of ISI counts (5 ms bins, 1 ms steps), normalized by number of trials and number of steps per bin (i.e. 5), show enhanced gamma band firing in γnsFS. The *vertical gray lines* demarcate the gamma-band (30 – 55 Hz, ISIs 18 – 33 ms; Wilcoxon signed rank test, * p<0.05). **G.** Median and Inter-Quartile Range (IQR) for each FS subtype for CV^2^ (*left*), ISI mean (*center*), and ISI variance (*right*; Wilcoxon signed rank test for hit miss comparisons, * p<0.05). **H.** ISI peak distribution on hit and miss trials. On hit trials, the plnsFS showed a significant rightward shift, driven by the emergence of a subgroup expressing peak gamma band ISIs (*right*; Wilcoxon signed rank test, * p<0.05).

**Figure 2.**
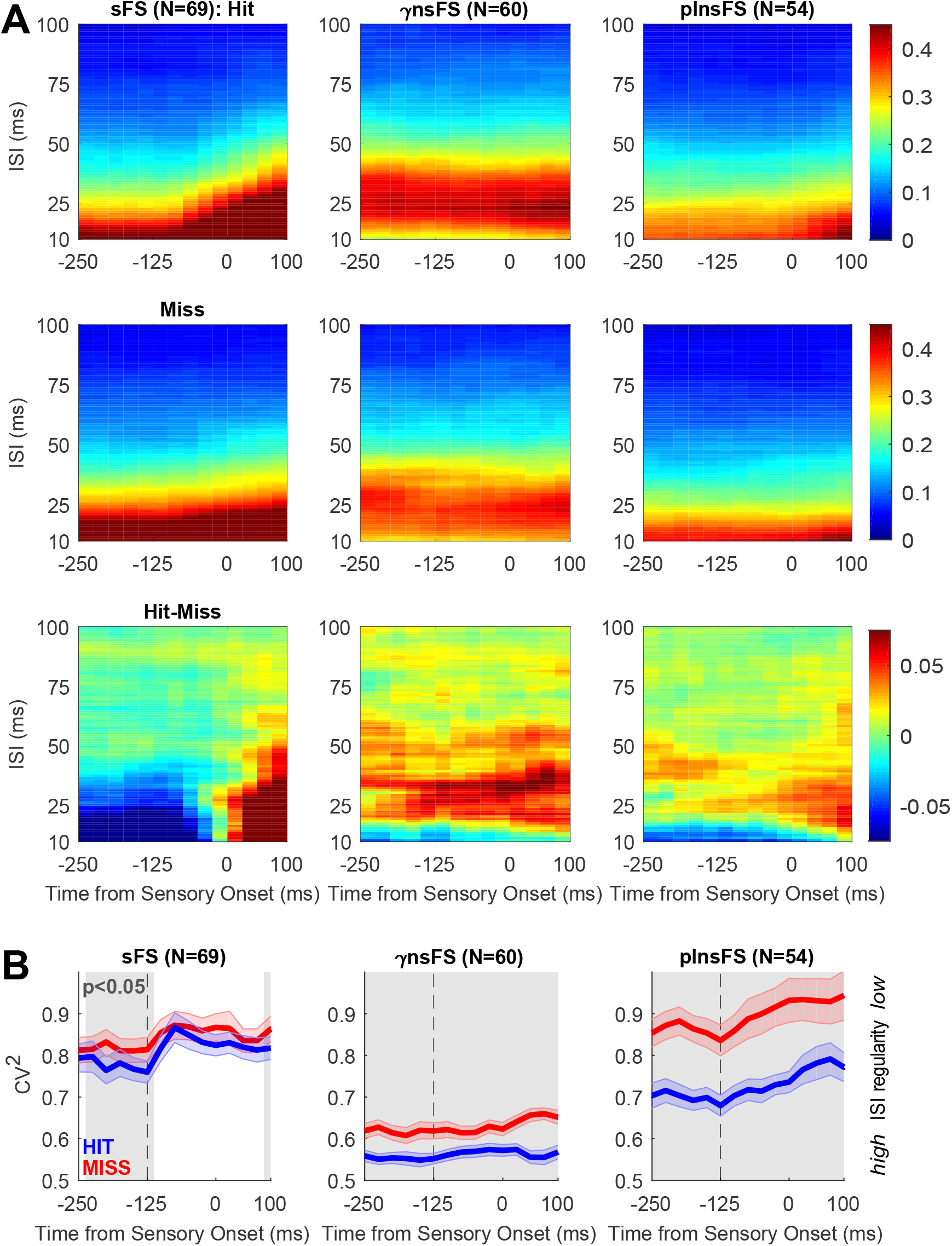
γnsFS Show Enhanced Gamma Interval Spiking on Hit Trials Before and After Sensory Onset. **A-B.** Normalized ISI histograms and CV^2^ in 250 ms – long time windows, sliding in 25 ms steps. X-axis denotes the center of the 250ms time window. Sensory onset is included in analysis window starting X=-125 ms. **A.** Normalized ISI histograms are mapped over time on hit trials (*top*), misses (*middle*), and hits-misses (*bottom*). Color-scale is fixed for each row (*right*). Enhanced gamma interval spiking that predicts successful sensory performance is evident in γnsFS beginning before stimulus onset and continuing poststimulus, whereas plnsFS showed emergence of gamma spiking poststimulus. For sFS, enhanced firing poststimulus is not specific to the gamma-band, but reflects a generalized increase in rates (see **Supplemental Figure 1E**). **B.** Non-sensory responsive FS of both subtypes showed more regular firing on hit trials before and after sensory onset (mean CV^2^ ± SEM on hits *blue* and misses *red). Gray* shading indicates a significant difference between hits and misses, pointwise Wilcoxon signed rank test p<0.05).

The γnsFS showed a different pattern. This subtype was identified by its gamma-range ISI distribution, and accordingly showed a band of higher probability ISIs at ~25 ms (40 Hz) that was persistent until the reaction time, spanning the period of sensory onset (**Figure 2A, *middle column***). While gamma interval spiking was high in this group on both hits and misses, activity in the gamma ISI band was higher on hits than misses for all time points from the prestimulus period until 100 ms after sensory onset. This finding indicates that increased gamma spiking in γnsFS was present until the sensory decision was relayed. The plnsFS showed a pattern similar to γnsFS when the difference in ISIs was mapped between hits and misses. The spectral power of spike trains shows a similar effect to these inverse ISI distributions, as expected (**Supplemental Figure 3A**).

We next asked whether increased ISI regularity (lower CV^2^) on hit trials, observed in all 3 subtypes prestimulus, persisted into the poststimulus period. As in **Figure 2A**, we applied a 250 ms sliding window. The shorter 250 ms window, compared to the 1 s window used in **Figure 1**, leads to a lower CV^2^ absolute value. However, the rank order of CV^2^ values was preserved across FS using either measurement window (**Supplemental Figure 3B**). In addition, baseline CV^2^ values did not depend on baseline rate, indicating that CV^2^ differences are not a byproduct of rate differences (**Supplemental Figure 3C**).

The sFS showed a decrease in regularity, i.e. an increase in CV^2^, at sensory onset, consistent with the interruption in internally generated spiking by sensory drive in the sensory responsive group. Poststimulus, the differences in regularity observed prestimulus were lost, consistent with the increase in firing rate in this group after vibrissa deflection (**Figure 2B *left*.** Note that sensory onset is included in the 250ms sliding window starting X=-125 ms). In contrast, the lower CV^2^ among γnsFS compared to the other groups (described in **Figure 1C**) persisted into the poststimulus period, on hit and miss trials (**Figure 2B *center***). In further contrast to sFS, both nsFS subtypes showed significant differences in CV^2^ in the peristimulus period that were significant for all time points tested, between −1000 ms prestimulus to 100 ms poststimulus, i.e. until the reaction time.

Fano Factor quantifies the trial-by-trial variability in firing rate. A renewal process is defined as having independent and identically distributed ISIs, with no effect of history in consecutive ISIs. Accordingly, a renewal process is characterized by identical CV^2^ and Fano Factor values. In the prestimulus period, CV^2^ and Fano Factor were positively correlated across all FS, with the sFS subtype having a lower correlation than the two nsFS subtypes (**Supplemental Figure 3D**; Pearson’s correlation across all trials: sFS R=0.387, 95% CI 0.166 – 0.572; γnsFS R=0.702, CI 0.546 – 0.811; plnsFS R=0.740, CI 0.588 – 0.841). However, the slope of the CV^2^ and Fano Factor linear regression is not at the identity line, suggesting that FS spike generation is not a renewal process (ratio of Fano Factor to CV^2^ across sFS has a median of 2.31 and IQR 2.01 – 2.81; γnsFS median=2.21, IQR 1.80 – 2.52; plnsFS median=1.96, IQR 1.68 – 2.70). This difference is consistent with state dependencies in spike processes, which manifests in part as hit and miss differences in our data.

After sensory onset, Fano Factor decreased on hits and misses (**Supplemental Figure 3E**), as previously reported (Churchland et al., 2010). In contrast, CV^2^ increased after stimulus onset. These results suggest that sensory drive to sFS causes a transient decrease in sFS spike count variability across trials (decrease in Fano Factor), while simultaneously increasing CV^2^.

In sum, sensory onset led to significant changes in various metrics including rate, CV^2^, Fano Factor, peak ISI and spike gamma-band power in the sFS subtype. In contrast, none of these metrics changed at sensory onset in γnsFS (**Supplemental Figure 3F**). These results indicate that mechanisms that underlie γnsFS spike generation are not perturbed by sensory drive.

### Rhythmic Gamma Spiking by γnsFS is Not Reset by Sensory Stimulation on Detected Trials

The ISI heat maps, and related statistical measures, indicate constancy in the firing patterns of γnsFS from pre-to poststimulus (**Figure 2, Supplemental Figure 3B**). However, in a rhythmic process, an external perturbation could reset the phase of an ongoing gamma oscillation, even in the absence of gross changes from pre-to poststimulus. Reset due to sensory drive would affect the interval between the last prestimulus spike and the first poststimulus spike, referred to here as the *peristimulus* ISI. The *last prestimulus* ISI refers to the prestimulus ISI immediately preceding the peristimulus ISI (**Figure 3A**).

**Figure 3.**
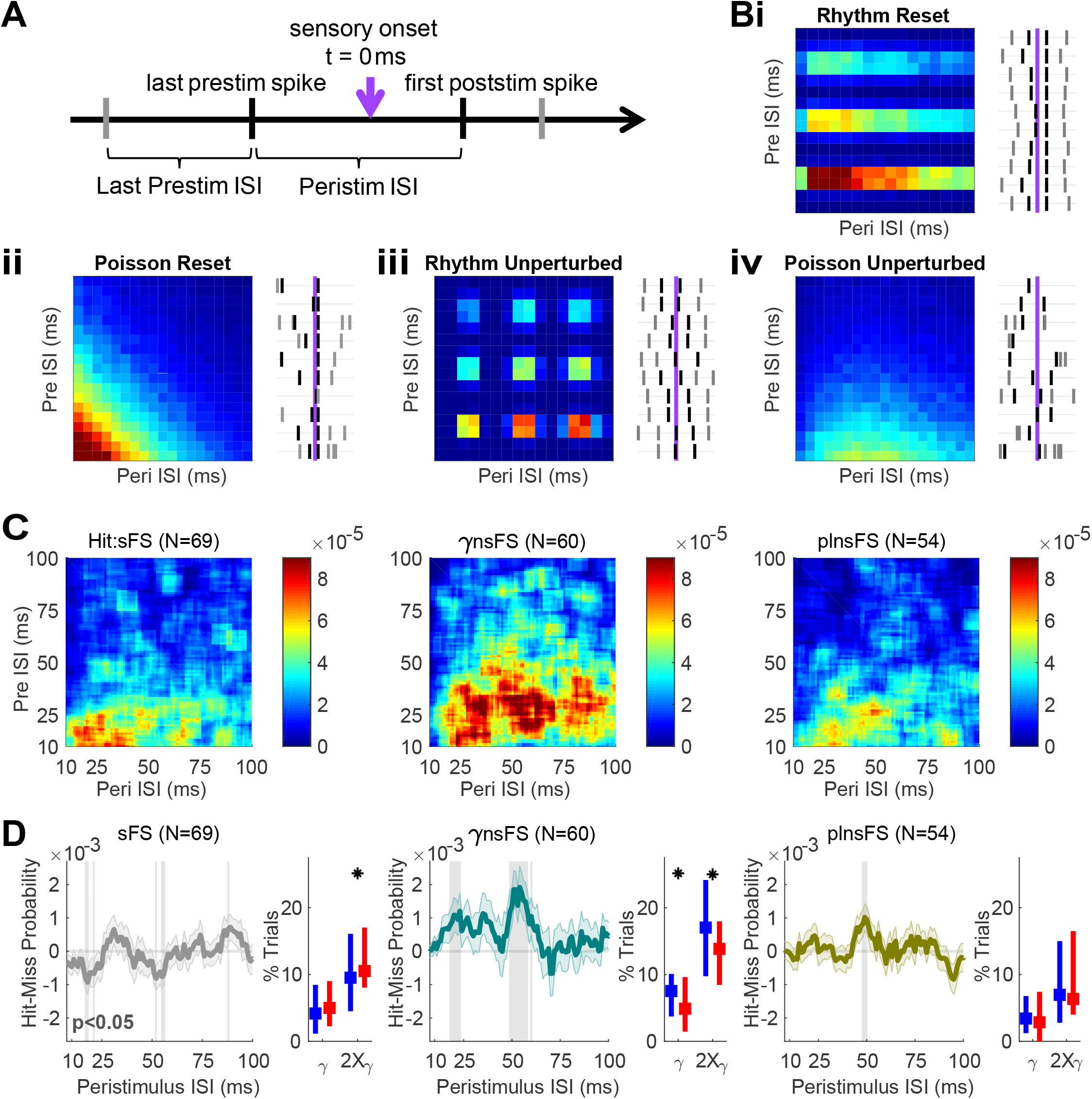
Gamma Interval Spiking in γnsFS is Not Reset by Sensory Onset on Detected Trials. **A.** As a test of whether prestimulus spiking persisted into the poststimulus period, we calculated the relationship between the last *prestimulus* spike interval before sensory onset and the interval spanning sensory onset (the *‘peristimulus’* interval). **B.** Simulations: To better understand the potential impact of sensory input on spike probability, we simulated four alternative spike generation processe (see *Methods* for details). (i) Gamma rhythmic prestimulus spiking ‘reset’ by a sensory input that realigned the first poststimulus spike with a fixed latency from sensory onset; (ii) Poisson spike generation prestimulus ‘reset’ by sensory onset, driving responses at a fixed latency (iii) Gamma rhythmic prestimulus spiking that was unperturbed by sensory onset; and, (iv) Poisson spiking that was unperturbed. Data are plotted as a two-dimensional map of spike probabilities with the peristimulus ISI on X-axis, and the last prestimulus ISI on Y-axis (*left*). For each simulation, 10 examples of the simulated spike ratergrams are shown (*right*). Simulation parameters were chosen to result in a mean firing rate of 10 Hz, to match the baseline firing rates observed across FS subtypes (**Supplemental Figure 1**). For the rhythmic spiking process, this was achieved by randomly thinning 3 out of 4 spikes generated from a gamma rhythmic process (peak ISI of ~25 ms). This period ‘skipping’ is reflected as high density at integer multiples of the peak ISI, i.e. 50 and 75 ms. **C.** Data: 2D normalized ISI histograms (10 ms bins, 1ms steps on either axis). The ISI count was normalized by number of trials and number of steps per bin (i.e. 10×10=100). Color reflects the mean firing rate on hit trials (fixed color-scale across FS subtypes, *right*). **D.** Differences in peristimulus interval probabilities on hit-miss trials (i; mean and SEM, 10 ms bins, 1 ms steps; pointwise Wilcoxon signed rank test, * p<0.05). *Gray* shaded regions indicate significant differences on hits compared to misses. (ii) The median and IQR of the % of trials with peristimulus ISIs in the gamma range (18 – 33 ms) or at its period doubling (36 – 67 ms; Wilcoxon signed rank test, * p<0.05).

To test the possibility of reset upon sensory onset in γnsFS gamma rhythmic spiking, we looked at the relationship between the last prestimulus ISI and the peristimulus ISI. Several possible outcomes of this analysis were modeled (diagrammed in **Figure 3B**). We simulated two possible spike generation mechanisms, one rhythmic (**Figure 3Bi, iii**), and one Poisson (**Figure 3Bii, iv**). For each simulated spike train, we compared the case where the first poststimulus spike occurred at a fixed latency to sensory onset (e.g., was evoked or reset by sensory drive; **Figure 3Bi, ii**), versus the case where sensory onset had no effect on the spike train (**Figure 3Biii, iv**). The relationship between the last prestimulus ISI and the peristimulus ISI was visualized as 2D normalized histograms in the simulations and data, where the X-axis is the last prestimulus ISI and Y-axis the peristimulus ISI. The X-projection of this 2D histogram corresponds to the normalized histogram for the last prestimulus ISI, while the Y-projection corresponds to the normalized histogram for the peristimulus ISI.

The prestimulus ISI histogram would be expected to be exponential for Poisson spike trains. For the peristimulus ISI in a sustained Poisson process (**Figure 3Biv**), the implementation of uniformly random inter-trial intervals leads to an elongation of peristimulus ISI relative to the last prestimulus ISI (*Methods*). In a Poisson process with reset (**Figure 3Bii**), the peristimulus ISI distribution is also exponential, with a rightward shift that corresponds to the latency to the first poststimulus spike (set to 12.5 ms in the simulation). In a rhythmic spike process where the sensory stimulus posed no interruption to an ongoing oscillation (**Figure 3Biii**), the ISI immediately prior to sensory onset (e.g. 25 ms) should predict another such period (e.g. a 40 Hz cycle), or if that period was skipped, fire at integer multiples of this period (e.g. at ~50 and ~75 ms ISI). Given that mean firing rates of γnsFS is ~10 Hz rather than 40 Hz, such period skipping is predicted, and was implemented in the model by randomly thinning 3 out of 4 spikes in a 40 Hz rhythmic spike train. In the case of rhythm ‘reset’ (**Figure 3Bi**), the peristimulus ISI distribution would be an exponential distribution with a rightward shift that corresponds to the latency to first poststimulus spike, as with the Poisson process with reset.

The 2D histograms of the three FS subtypes are shown in **Figure 3C**. The 2D histogram of sFS on hit trials (**Figure 3C *left***) paralleled the results predicted from a simulated Poisson spike train with reliable latency following sensory onset (**Figure 3Bii**). This result is expected, as the prestimulus ISI distribution of sFS is Poisson-like with a spike refractory period (**Figure 2A**), and a reliable latency to the first sensory evoked spike is a robust feature of sFS in barrel cortex (Simons and Carvell, 1989; **Supplemental Figure 4A-B**). In comparison, the plnsFS 2D histogram (**Figure 3C *right***) resembled the model of an uninterrupted Poisson process (**Figure 3Biv**). The 2D histogram of the γnsFS (**Figure 3C *center***) showed 3 distinct peaks in peristimulus ISI density at ~25, 50 and 75 ms. This periodicity closely mirrors the model prediction for an unperturbed gamma oscillation with random spike omissions (**Figure 3Biii**), as opposed to a rhythm that is reset upon sensory onset (**Figure 3Bi**). The lack of reset in γnsFS is further supported by the absence of a peak in the latency to the first poststimulus spike (**Supplemental Figure 4A-B**).

Next, we plotted the difference in ISI probabilities on hits versus misses (**Figure 3D**). The γnsFS showed rhythmic enhancement on hits relative to misses, with more ISIs at ~25 and ~50 ms. These effects were present but less prominent in the plnsFS. The sFS exhibited the opposite behavior, with significant decreases in the 25 and 50 ms ranges. In sum, gamma rhythmic spiking of γnsFS is not reset by sensory drive, and this sustained, regular and periodic spiking at gamma in γnsFS predicts higher detection task performance.

### Gamma Spiking by γnsFS is Negatively Correlated with the Gamma LFP, and the Gamma LFP Negatively Predicted Detection

The role of gamma in perception is often assessed by measuring LFP power, and/or spiking coherence with LFP in the gamma-band. Several cogent critiques of the appropriateness of gamma for binding hypotheses are, for example, based on this measure. In the present study, gamma-band LFP power (30 – 55 Hz) was lower on hits compared to misses pre- and peristimulus, as evident in the mean power spectral densities for hits and misses (**Figure 4A, Supplemental Figure 4C** for peristimulus spectrogram). The LFP gamma power in the prestimulus period (−250 – 0 ms) was higher on misses than hits throughout the range of stimulus amplitudes. On catch trials, the prestimulus LFP gamma power was higher on false alarms than correct rejections (**Fig 4B**). This negative relationship between the prestimulus LFP gamma power and detection performance is also clear when looking at hit rate in quintiles of gamma-band power (**Fig 4C**).

**Figure 4.**
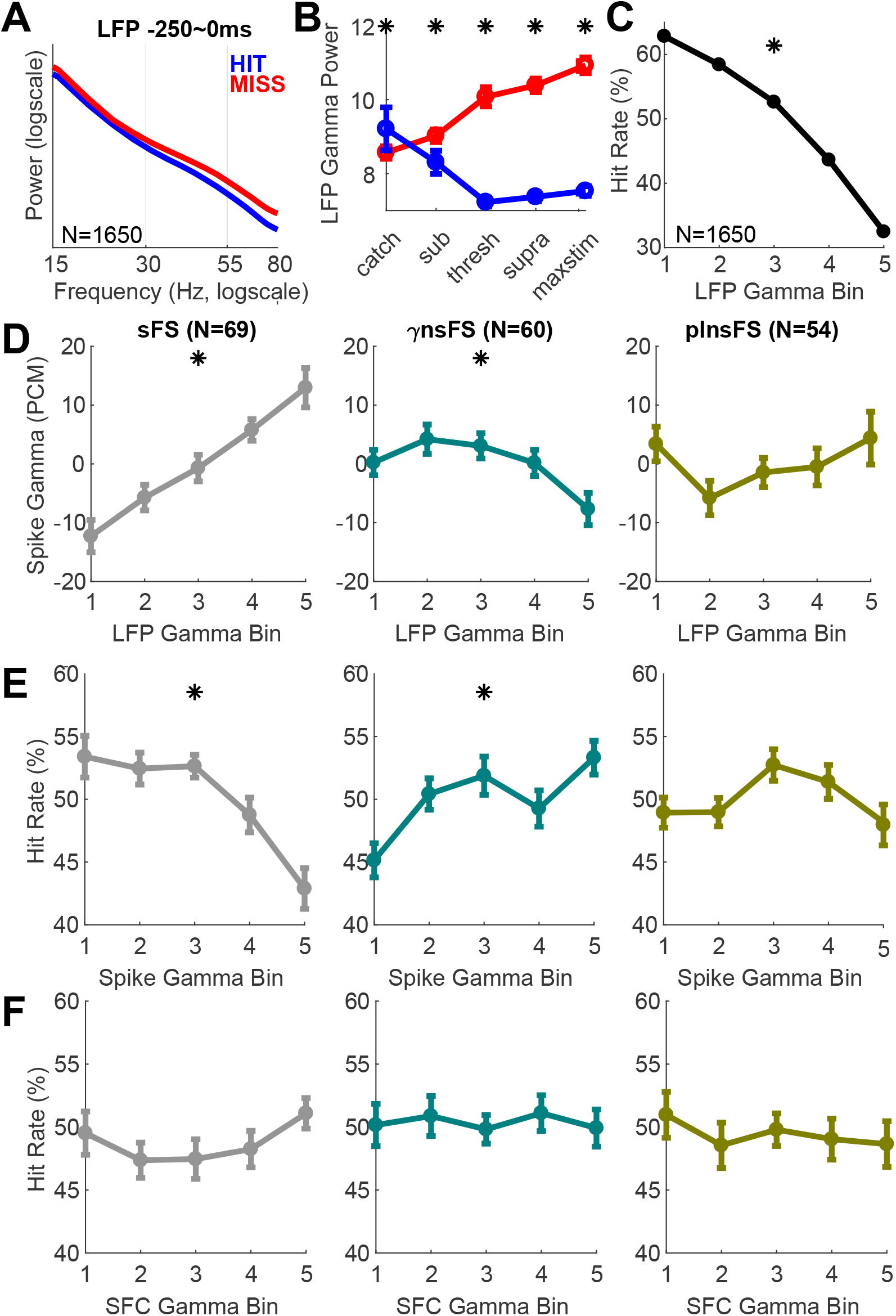
Spiking Rhythmicity in Non-Sensory FS is not Reflected in the Gamma Band LFP: Sensory FS Spike Gamma Power and the LFP Gamma Power Predict Unsuccessful Performance. **A.** LFP power spectral density calculated −250 – 0 ms prestimulus on hits (*blue*) and misses (*red*; Mean ± SEM across tetrodes, N=1650). **B.** Same as **Supplemental Figure 1G**, but for the LFP power in the gamma-band (30 – 55 Hz), prestimulus (−250 – 0 ms). **C-F.** Relation between the gamma-band LFP power, spike power, SFC and hit rate, in the −250 – 0 ms prestimulus period. Stimulus amplitude matched trials (hits and misses pooled) were sorted into quintiles, in ascending order from 1 to 5 (*left* to *right*). The LFP measurements were taken from adjacent electrodes to those recording the spikes. **C.** Hit rate as a function of LFP gamma power. Mean ± SEM hit rate across tetrodes (N=1650) is plotted for each quintile. Wilcoxon signed rank test was applied to test whether LFP gamma power averaged across hits and misses were significantly different across tetrodes (* p<0.05). **D.** Gamma-band power for spikes, calculated as percent change from the mean (PCM) and plotted as a function of LFP gamma power (mean ± SEM). In addition, for each FS, the trial-by-trial Spearman correlation coefficient was calculated between LFP gamma band power and spike gamma power. Wilcoxon signed rank test was applied to determine whether the distribution of Spearman correlation coefficients across each FS subpopulation was significantly different from 0 (* p<0.05). **E.** Same as **C**, but for gamma-band power of multitapered Fourier transformed spike trains. Mean ± SEM hit rate across each FS subtype. **F.** Same as **C** and **E**, but for gamma-band SFC across each FS subtype. Across the entire stimulus amplitude range, the LFP gamma power was lower on hits than misses. On catch trials, false alarms had a significantly higher LFP gamma power than correct rejections.

To assess the relationship between spike spectral power (**Supplemental Figure 3Aii**) and LFP spectral power, we looked at the trial-by-trial correlation between these two quantities in the gamma-band (30 – 55 Hz; **Figure 4C**). The correlation between spike and LFP gamma-band power is akin to the often measured spike-field coherence metric (SFC), with the major difference being that the effect of phase consistency is not incorporated. In addition, zero spike trials are ignored when calculating SFC, but contributes to the correlation in our analysis. To prevent spike contamination of LFP from influencing our results (Ray and Maunsell, 2011; Waldert et al., 2013), we calculated the spike – LFP correlation in gamma-band power by using the LFP averaged from three tetrodes neighboring the tetrode where the spikes were recorded from.

Gamma-band LFP power was correlated with spike spectral power in the gamma-band among sFS (Figure 4C *left*, Spearman correlations are significantly greater than zero; p=5.66χ10^−9^ one-sample Wilcoxon signed rank test). In contrast, higher gamma spiking power in γnsFS was negatively correlated to LFP gamma power (**Figure 4C *center*** p=8.99χ10^−5^). The plnsFS had no significant relationship to LFP gamma power (Figure 4C *right*,p=0.377). These relationships between gamma-band LFP and spiking were preserved poststimulus before reaction time (**Supplemental Figure 4D**, 0 – 100 ms period; sFS p=2.13χ10^−9^; γnsFS p=0.00159; plnsFS p=0.708 Wilcoxon signed rank test).

Consistent with the prestimulus portion of the hit minus miss spike spectrogram in **Supplemental Figure 3Aii**, the hit rate for quintiles of gamma-band power spiking shows a negative relationship for sFS (**Figure 4D *left***, p=2.46×10^−5^ Wilcoxon signed rank test), and a positive relationship for γnsFS (**Figure 4D *center*** p=0.0030). The plnsFS spike gamma power was not different between hits and misses (**Figure 4D *right***, p=0.382).

Unlike spike spectral power and LFP power in the gamma-band, SFC in the gamma-band did not differ between hit and miss trials, in any of the FS subtypes (**Figure 4E**; sFS p=0.163; γnsFS p=0.354; plnsFS p=0.126 Wilcoxon signed rank test).

Next, we tested whether the LFP and spike gamma power in the poststimulus period depended on the stimulus amplitude. Ray and Maunsell (2010) pointed out that dependence of the LFP gamma on specific stimulus features undermines its potential role as a binding template. In agreement, the LFP gamma power increased monotonically with increasing stimulus amplitude (**Supplemental Figure 4E**). The sFS spike gamma power also linearly depended on stimulus amplitude, but γnsFS and plnsFS spike gamma power did not, adding support to their independence from specific stimulus features (**Supplemental Figure 4F**). Note, the spike spectral power generally correlated positively with firing rate on a trial-by-trial basis, reflected in the similarity between **Supplemental Figure 4F** and **Supplemental Figure 1G**.

In sum, these findings are consistent with prior claims (Pritchett et al., 2015; Ray and Maunsell, 2015) that gamma band LFP power does not index a viable temporal carrying signal that promotes sensory processing. However, persistent spiking gamma observed in the γnsFS predicted perceptual success, was not perturbed by sensory input, and was negatively related to the gamma band LFP. As such, this predictive temporal pattern in spiking data could reflect a coordinating mechanism that could potentially enhance binding or communication-through-coherence.

## Discussion

The present study shows that persistent gamma spiking among non-sensory FS predicts perceptual success. These effects are most pronounced in a subtype (γnsFS) whose persistent gamma activity positively predicted performance, and whose spiking was negatively related to the gamma LFP. As such, γnsFS could, in concept, facilitate the temporal organization of local neocortical firing amongst an informative ensemble of cells. Importantly, γnsFS spiking is free of the stimulus dependence of the LFP gamma that, to date, has provided a strong argument against theories proposing a temporally-coordinating role for gamma.

Given that non-sensory FS represented the majority of FS encountered, they are positioned to play a substantial role in the coordination of local brain activity. Prior studies, including our own, that have tested the association between neocortical gamma and behavioral performance have focused on the relationship between spiking and the LFP, and have not analyzed endogenous FS spiking periodicity as an independent variable. More generally, when spiking patterns are analyzed, sensory responsive neurons are typically the focus. Exclusive reliance on either of these common analysis choices would prevent observation of the functionally defined FS subtypes reported here.

The present study is focused on the initial description of these distinct FS subtypes during behavior, reflecting the widely-accepted importance of FS for sensory processing. Our current recording strategy (implanted tetrodes) facilitated recording during extensive psychophysical characterization. This approach is not optimal for identification of the laminar position of recordings, and we therefore refrained from speculating on layers of origin. Despite these caveats, we note that γnsFS and sFS did occur on the same tetrode in the same session on several occasions, arguing against a strong spatial segregation between types, including a strict laminar segregation.

In primary sensory areas of neocortex, much attention has been directed to the fast (feedforward) and nonspecific (broadly tuned) sensory responsive properties of FS (Andermann et al., 2004; Andermann and Moore, 2006; Cruikshank et al., 2007; Haider et al., 2013; Isaacson and Scanziani, 2011; Kerlin et al., 2010). The majority of FS in the present study obtained during behavioral performance are non-sensory. This sparsity in sensory responsiveness is often observed among pyramidal neurons in primary sensory areas of the neocortex (Kerr et al., 2003; O’Connor et al., 2010; Hromádka et al., 2008; Kwon et al., 2016; Olshausen and Field, 2004; Peron et al., 2015), and the present findings extend this result to FS during an SI-related behavior.

Two precedents in the literature are suggestive of the subtypes described here. Recording in frontal neocortex, Puig et al. (2008) found two distinct activity patterns of FS during neocortical up states. One FS subtype demonstrated ISI distributions with peaks in the gamma-band that discharged late within an up state. Such spiking behavior strongly resembles that of the γnsFS described here in SI. Li and Huntsman (2014) reported two broad electrophysiological categories of FS within layer 4 of SI barrel cortex based on the onset of the first action potential in a depolarizing train. They found that Fs that showed delayed spiking in response to current injection also had smaller thalamocortical-evoked responses compared to early spiking FS, potentially reflecting the sensory and non-sensory subtypes observed in the present study.

The present data also imply different synaptic connectivity patterns of the three functionally-defined FS subtypes. Conclusive identification of these connectivity patterns would require direct manipulation of identified neurons exclusively within a subtype and/or anatomical reconstruction of identified FS connectivity with synapse-level precision. That said, the physiological characterization of these subtypes suggests their possible targets. The positive correlation between sFS and LFP suggests that sFS are a primary regulator of coordinated current fluctuations in cells with large dendrites that are the primary driver of the neocortical LFP (Buzsáki et al., 2012). Conversely, the negative correlation between γnsFS and LFP may reflect the targeting by this subtype of other interneurons. Consistent with this prediction, the ISI peak shift in plnsFS towards the gamma range on hits relative to misses may be the result of its entrainment by enhanced rhythmic spiking in γnsFS. Another possibility is that sFS and γnsFS are mutually inhibitory, reflecting their opposite relationships to gamma-band LFP power. However, the total lack of influence on γnsFS spiking by sensory drive suggests a decoupling of inputs from the sensory-responsive units of the local network to the γnsFS.

A potentially important implication of these results is that a distinct subtype of interneurons, that acts independently of the sensory processing, is the carrier of perceptually-relevant oscillations. Testing the hypothesis that persistent gamma rhythmic spiking among non-sensory FS would enhance perception, through binding or communication-by-coherence, requires further evidence (Singer, 1993; Singer and Gray, 1995). First, the hypothesized role of persistent rhythmic spiking as a coordinating ‘reference’ signal, present before and after arrival of the information it coordinates, could in part be tested by examining the effect of peristimulus persistence on perception, compared with the effect of prestimulus alone and poststimulus alone. Second, testing the hypothesis that rhythmic spiking of γnsFS serves a role in binding across multiple brain areas requires the demonstration of such neurons outside of SI, and recording and controlling their coherence at multiple sites while the experimental subject performs a task requiring feature integration. If supported by investigation in other paradigms and brain areas, our findings provide a unique resolution to a consistent challenge to theories predicting a role for oscillations in optimal brain communication.

## Methods

### Animals

Electrophysiology was collected from four mice (3 males, 1 female) performing a vibrissae deflection detection task. Mice were 8-15 weeks at the time of surgery, and were recorded from for up to 7 months. Animals were individually housed with enrichment toys and maintained on a 12 hr reversed light-dark cycle. All experimental procedures and animal care protocols were approved by Brown University Institutional Animal Care and Use Committees and were in accordance with US National Institutes of Health guidelines.

### Driveable Electrode Implants

We implanted flexDrives loaded with 16 tetrodes and constructed in house (Voigts et al., 2013). Tetrodes were made with Sandvik Kanthal HP Reid Precision Fine Tetrode Wire, Nickel-Chrome 0.012 mm diameter, and gold-plated to 200-400 kΩ impedence. The guide tube array was made with 33 ga polyamide tubes, resulting in ~250 μm spacings. Two stainless-steel screws (0.6 or 0.8 mm diameter, 0.5 or 1 mm length) were electrically connected (soldered) to the electrode interface board (EIB) through stainless steel wire. These skull screws were implanted through skull to rest on top of the dura to serve as ground: one was positioned anterior to bregma and the other on the right hemisphere.

### Surgical Procedure

Details of the surgical procedure and behavior control was reported in a prior study (Shin et al., 2017). Briefly, mice were induced with isofluorane anesthesia (0.5 – 2 *%* in oxygen 1 L/min) and secured in a stereotaxic apparatus. We injected slow-release buprenorphine subcutaneously (0.1 mg/kg; as an analgesic) and dexamethasone intraperitoneally (IP, 4 mg/kg; to prevent tissue swelling). Hair was removed from the scalp with hair-removal cream, followed by scalp cleansing with iodine solution and alcohol. Then, the skull was exposed by scalp incision. After the skull was cleaned, muscle resection was performed on the left side. A titanium headpost was affixed to the skull with adhesive luting cement (C&B Metabond). Next, a ~ 1.5 mm–diameter craniotomy was drilled over barrel cortex of the left hemisphere, and subsequently a durotomy was performed. The guide tube array was centered at 1.25 mm posterior to bregma and 3.25 mm lateral to the midline. The drive body was angled 30 degrees relative to vertical to compensate for the curvature of barrel cortex. Once the implant was stably positioned, C&B Metabond and dental acrylic (All for Dentist) was placed around its base to seal its place. A drop of surgical lubricant (Surgilube) prevented dental acrylic from contacting the cortical surface. Mice were given ≥3 days to recover before the start of water restriction.

### Recording and Preprocessing

All electrophysiology data were collected using the Open Ephys system continuously, with a sampling rate of 30 kHz. Trial alignment was achieved through a synchronizing pulse output from the computer running the behavior control software, via an analog channel of a NI-DAQ device.

Local field potentials (LFP) were defined as the continuous data down-sampled to 1000 Hz. Median filtering was applied concurrent with down-sampling, i.e. down-sampling was conducted by choosing the median every 30 samples. Spike detection was conducted online during data acquisition to visualize the relevance of a recording site, with the online criterion for spikes was whether 300 – 6000 Hz bandpass filtered data crossed a permissive threshold of −50 μV, in at least one of the four electrodes comprising each tetrode. Offline, spike threshold for each electrode was readjusted as 4 times the standard deviation of 300 – 6000 Hz bandpass filtered data. The standard deviation was approximated as the median value divided by 0.6745 (Quiroga et al., 2004). Multi-unit activity (MUA) for each tetrode was defined as spikes that crossed this threshold in at least one of the four electrodes of that tetrode (one MUA per tetrode, per session).

To obtain single unit activity (SUA), offline spike sorting was conducted manually using Simple Clust (www.github.com/open-ephys/simpleclust). Only the clusters that were well isolated were classified as SUA. After sorting, single units were classified into regular-spiking (RS) and fast-spiking (FS) units, based on the time between peak to trough (T_PT_) in their average spike waveform (RS if T_PT_>0.4 ms, FS if T_PT_≤0.4 ms; Supplemental Figure 1D).

Tetrodes were lowered at the end of a session if the experimenter noticed that there were no single units detected from online sorting. Tetrodes were lowered by 1/8 turns, corresponding to ~31.25 μm.

### Behavior Training

Mice were water restricted for ≥7 days before start of training, during which time they were acclimated to being head-fixed on a fixed-axis styrofoam ball, where they were allowed to run freely. Water was delivered through a syringe after each acclimation session.

When training began, the vibrissae were secured through a suture loop. All macrovibrissae posterior to the 4^th^ arc on the right side were secured ~3 mm from the mystacial pad. The suture loop, in turn, was fed through a glass capillary tube (0.8 mm outer diameter) glued to a piezoelectric bender (Noliac CMBP09). On each trial, 20 Hz vibratory vibrissae stimulus train (10 deflections/500 ms) were delivered through the piezoelectric bender in the caudorostral direction, with a half-sine wave velocity profile that had fast rising phase (6 ms) and a slower relaxation phase (20 ms). The 10 deflections within each stimulus train of a trial were of the same amplitude. On a trial-by-trial basis, the stimulus amplitude was varied between 0 (a catch trial) to maximal amplitude (~1 mm deflection) in a randomized manner. Before the start of each session, the experimenter set the overall percentage of maximal trials and 0 amplitude catch trials. The rest of the trials were submaximal trials, where the stimulus amplitude was randomly drawn from a uniform distribution between 0 and maximal amplitude. Throughout training, the percentage of maximal amplitude trials was gradually lowered to 10%, and the percentage of catch trials was gradually increased to 25%.

The mice initially learned the vibrissae deflection to reward association in sessions where every vibrissae deflection trial was paired with ~3 μl water delivery regardless of the animal’s response. Vacuum suction followed 500 ms after every reward delivery, which removed any excess water not consumed by the animal in that trial. Mice learned the vibrissae deflection to reward association after about a week of reward-all training, at which point the animals would lick in anticipation of the reward with a stereotyped reaction time of ~250 ms poststimulus onset (median 254 ms; IQR 187 – 389; **Supplemental Figure 1C**). When this behavior was observed (>4 days of training), the reward-all phase of training was concluded.

During the subsequent phase performance was required for reward, and it was only delivered on detected (hit) trials where the mouse correctly reported detection by licking within 700 ms of the stimulus onset (reward window). Here, the consequence of non-detection (misses) was omission of reward. If the mouse falsely reported detection by licking during the 700 ms catch window, the trial outcome was classified as a false alarm. The consequence of false alarms was a 15 s timeout from the task, effectively delaying the next opportunity to obtain a reward. If the mouse correctly refrained from licking during this catch window, the outcome was a correct rejection. Inter-trial intervals (ITIs, defined as the interval between reward/catch window onsets) were randomly drawn from a uniform distribution ranging between 4.5 – 8 s. There was no cue indicating the start of each trial.

If the animal developed a habit of excessive impulsive licking, an ITI-reset was implemented until that habit was eliminated. In these sessions, licking during the ITI would prolong the ITI up to 10 times.

Mice were weighed before and after each session. If the mouse consumed ≥1 ml of water throughout a behavior session, the mouse would gain ~0.7g at the end of a behavior session. Hence, if the mouse did not gain ≥0.7g during the behavior session, supplementary water was given several hours after the conclusion of the session, so that the mouse would have drunk at least 1 ml each day. In addition, mouse weight was monitored throughout the duration of training and maintained ≥80% of their original weight at the time of surgery. The duration of training ranged between 3 to 7 months.

### Behavior Analysis

A typical behavior session lasted ~2.5 hours, with ~1000 trials (median 946, IQR 817 – 1032). Even in well trained mice displaying robust psychometric behavior and stereotyped reaction time, there were periods where animals defaulted to non-optimal strategies, such as excessive impulsive licking or non-engagement. All hit, miss, false alarm and correct rejection trials analyzed were trials where there was no licking activity up to 1 s before the onset of the reward/catch window. Across sessions, the percent of trials filtered out due to licking in the 1 s prestimulus had a median of 10 %, with IQR 7 – 26 %.

In addition, we employed d’≥1 criteria to filter out periods of low performance. The d’ was calculated in sliding windows of 51 trials advanced in 1 trial steps. For any blocks with d’≥1, the middle trial was included in the analysis. The first and last 25 trials were included if the first and last 51 trials had d’≥1, respectively. The d’ was defined as the following:

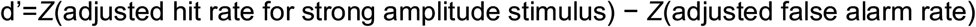

where *Z* stands for the inverse of the Gaussian distribution function. For each session, N strongest amplitude stimulus was analyzed for the *Z* score, where N equals the number of catch trials in that session. Hit rates and false alarm rates were adjusted to avoid infinite values of d’ by adding 0.5 to the number of trials in each category (Hautus, 1995; Macmillan and Creelman, 2004; Miller, 1996), i.e.:

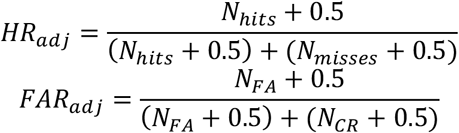

where *HR_adj_* denotes adjusted hit rate and *FAR_adj_* denotes adjusted false alarm rate; and *N_hits_,N_misses_,N_FA_,N_CR_* denotes the number of hits, misses, false alarms and correct rejection trials, respectively. The d’ filtering left a median of 30% trials, IQR 10 – 45 %. In many cases, this was due to satiation at the end of the session. As such, the percentage of d’<1 trials between the first d’≥1 trial of the session and the last d’≥1 trial of the session was much lower, with a median of 5% and IQR 0 – 22 %.

For each session, psychometric curves were calculated by sorting the d’≥1 trials in ascending stimulus amplitude, and smoothing the sorted stimulus amplitude vector and the corresponding trial outcome vector (1 for hits, 0 for misses) with 51 trial windows in 1-trial steps (**Supplemental Figure 1A**). Only the behavior sessions with a psychometric curve that had a good fit with a Boltzmann distribution were selected for analysis (across the 128 sessions, R-squared median 0.96, IQR 0.93 – 0.98; root mean squared error (RMSE) median 0.067, IQR 0.053 – 0.086).

To compare hit and miss trials independent of variations in stimulus amplitude, we performed for each session stimulus amplitude histogram matching of hit and miss trials, such that matched hits and misses were equal in trial count and stimulus amplitude distribution. Matching was conducted on submaximal trials with d’≥1. In addition, hit trials were limited to trials with ≥100 ms reaction time, such that analysis of stimulus evoked activity between 0 to 100 ms from stimulus onset would be free of motor activity associated with licking. The application of the RT≥100 ms criteria among d’≥1 submaximal hit trials yielded a median of 98% of preserved trials across sessions, with IQR 94 – 99 %. The histogram matching process was as follows (**Supplemental Figure 1B**):

1. Trials eligible for matching (submaximal, d’≥1, RT≥100ms) were divided into 15 bins, equidistant in stimulus amplitude.
2. For each bin, hits and misses were matched in number; if there were more hits than misses, we randomly sub-selected hit trials equal to the number of miss trials; and vice versa for bins with more misses than hits.

Across sessions, this resulted in a median of 78 matched hits and misses each per session, where IQR was 57 – 95 trials.

### Details of Data Analysis

Data analysis was performed in MATLAB (Mathworks). Sensory responsiveness was defined based on stimulus probability (SP, **Supplemental Figure 1E**). SP is defined as the area under the curve (AUC) for the receiver operating characteristic (ROC) curve in an ideal observer analysis, where the ideal observer discriminated maximal stimulus amplitude trials from zero amplitude catch trials based on firing rate in the 0 – 100 ms period poststimulus onset. 95% confidence interval of SP was determined by bootstrapping 1000 times. MATLAB function “perfcurve” was used for this analysis.

For each SUA and MUA recording, spike times were converted into spike trains in bin resolution of 1 ms (bin value is 1 if there was a spike at that time point, 0 otherwise). Spike trains were converted into matrices such that each column corresponded to a trial. Peristimulus time histograms (PSTH) and autocorrelograms were obtained from these matrices. The PSTH was calculated by 1) averaging across trials, 2) multiplying by 1000 such that the units would be in Hertz (Hz), then 3) smoothing with a Gaussian filter (with a standard deviation of 3 ms). Autocorrelograms were calculated by 1) truncating the spike train trial matrix to just the prestimulus period, 2) counting coincident spikes between the prestimulus matrix and the same matrix with offsets varying from −250 to 250 ms, 3) normalizing the number of coincident spikes by the total number of spikes in the original matrix, such that the resulting autocorrelogram would have a value of 1 at 0 offset.

The interval between two consecutive spike times of a SUA or a MUA is defined as the inter-spike interval (ISI). Coefficient of variance squared (CV^2^) is defined as the variance of the ISIs divided by the mean of ISI squared.

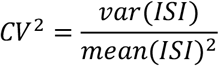

Lower CV^2^ implies higher regularity in ISI’s. ISI distribution and CV^2^ for a designated time window was calculated by first pooling across trials the ISIs contained within the designated period. Peristimulus ISI was calculated for each trial as the interval between the last spike before stimulus onset and the first spike after stimulus onset. Fano factor (FF) is defined as the variance over mean of the firing rate (FR) across trials.

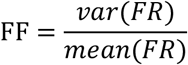

Spectral analysis for LFP and spike trains was calculated using a multitaper Fourier transform, where time-bandwidth product was set to 3 and the number of tapers was set to 5. Zero padding was applied such that the LFP or spike train data being processed would have a length of 2^N^; e.g. for a LFP segment with length of 250, 3 zeros were padded on either side such that the total length would be 2^8^=256. Chronux version 2.11 (http://chronux.org/, Mitra and Bokil, 2004) was used for this analysis. Spectral power in the gamma-band was calculated as the geometric mean across frequencies in the 30 – 55 Hz range.

For spike – LFP correlations presented in **Figure 4C**, the LFP was taken as the average of three tetrodes neighboring the tetrode where the FS spikes were recorded. This step was taken to prevent the influence of spike contamination on the LFP (Ray and Maunsell, 2011; Waldert et al., 2013).

### Spike Train Simulation

In **Figure 3B**, four spike trains were simulated: i) rhythm reset, ii) Poisson reset, iii) rhythm sustained and iv) Poisson sustained. For rhythmic spike trains (i, iii), ISIs were randomly drawn from a normal distribution with a mean of 25 ms and a standard deviation of 2.5 ms, then randomly thinned such that 1 out of 4 spikes would remain. The thinning was implemented to reflect γnsFS spike statistics, whose ISIs most commonly occurred in the gamma range (~25 ms), but showed a mean firing rate of ~10 Hz. For the Poisson spike trains (ii, iv), ISIs were randomly drawn from a exponential distribution with a mean of 100 ms (mean firing rate 10 Hz). Approximately 1.5×10^8^ ms spike trains were simulated, then divided into 10^5^ trials with randomized ITI uniformly distributed between 1 – 2 s. The implementation of uniformly distributed ITIs resulted in the elongation of the peristimulus ISI relative to the last prestimulus ISI in the unperturbed spike processes (iii, iv): For example, in a Poisson spike process, the exponential distribution of prestimulus ISIs (i.e. proportional to *e^−x^*; a Gamma distribution with a shape parameter of 1) becomes a distribution proportional to *xe^−x^* for the distribution of peristimulus ISIs (i.e. a Gamma distribution with a shape parameter of 2). For both spike processes, reset (i, ii) was simulated as fixed latency (12.5 ms) to spike after sensory onset, on every trial. The reset process results in the peristimulus ISIs being exponential distributed with a rightward shift that corresponds to the latency to first poststimulus spike.

## Acknowledgments

We thank Stephanie R. Jones, Scott J. Cruikshank, Barry W. Connors, Gabriela Manzano-Nieves, Manuel Gomez-Ramirez, Arif A. Hamid, Christopher A. Deister and Shai Sabbah for their comments on the manuscript. This study was supported by a grant from the US National Institutes of Neurological Disorders and Stroke to C.I.M. (R01NS045130), and fellowships from Fulbright and the Carney Institute for Brain Science to H.S.

---

**Supplemental Figure 1.**
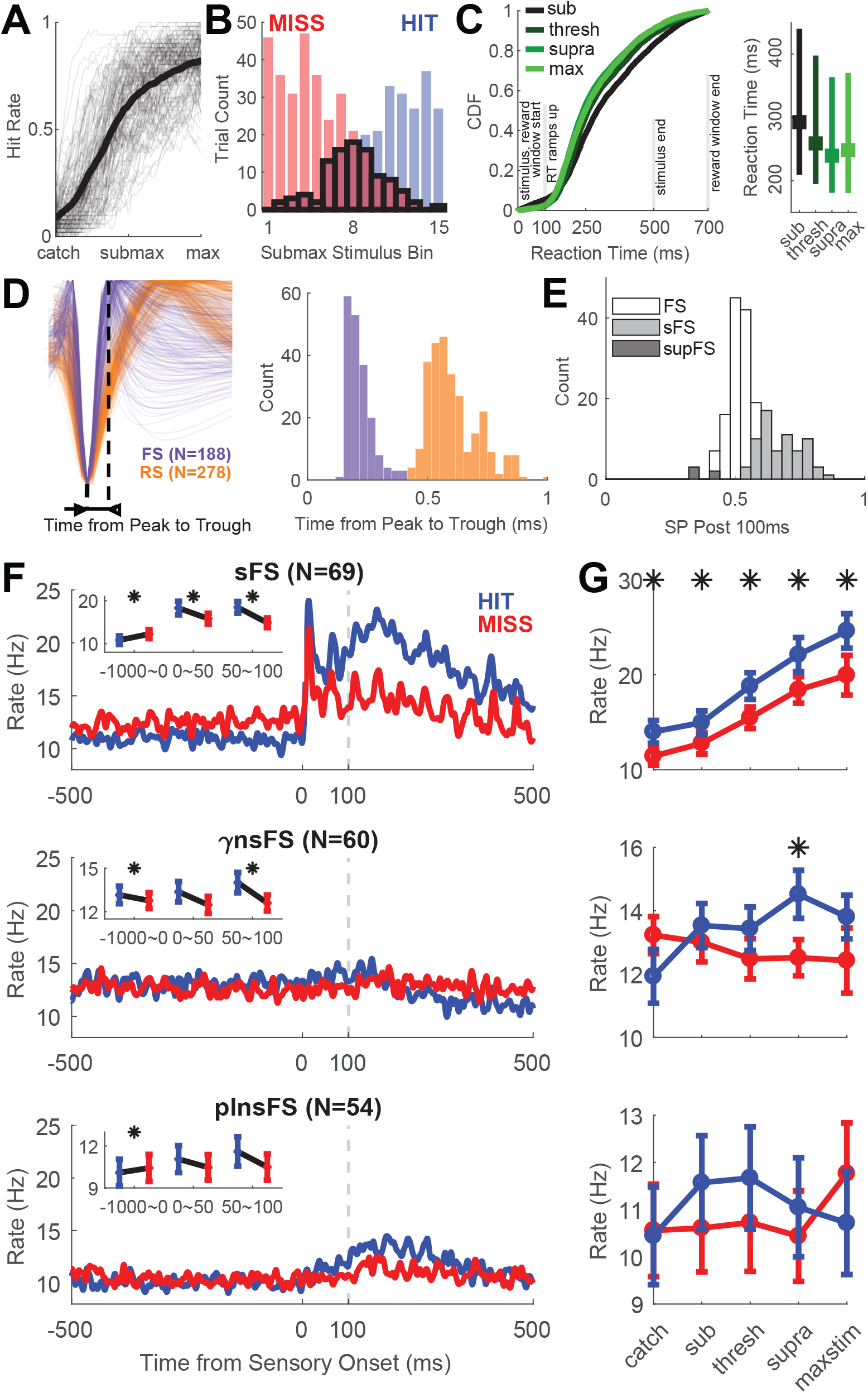
**A.** Psychometric behavior in a head-fixed vibrissae deflection detection task in every session (N=128, pooled across 4 mice, *gray lines* indicate each session, *black* the mean), showing increased hit rate as the sensory stimulus amplitude increased. On each trial, mice were trained to detect vibrissa deflections (10 deflections at equal amplitude, 500 ms, 20 Hz). Stimulus amplitude on each trial was randomly selected form a uniform distribution ranging from zero (catch trials) to maximal stimulation (~1 mm deflection). Hit trials were submaximal and maximal stimulus amplitude trials where the mouse reported detection within 700 ms of sensory stimulus onset (reward window). Licking during the reward window on catch trials resulted in a timeout of 15 s. Otherwise, inter-trial intervals (ITIs, intervals between reward/catch window onsets) were randomly drawn from a uniform distribution ranging from 4.5 – 8 s. To analyze only the periods where the animal was engaged in the task, we calculated d’ in sliding windows of 51 trials advanced in 1 trial steps, and filtered periods where the d’ < 1. For each session, psychometric curves were calculated from these d’ ≥ 1 trials. We selected 128 sessions from 4 mice displaying robust psychometric behavior. Psychometric curves from these sessions are plotted (*thin gray curves*): trials were sorted by stimulus amplitude, and both the sorted stimulus amplitude vector and the corresponding trial outcome vector (1 for hits, 0 for misses) was smoothed with a smoothing window of 51 trials. To calculate the grand average across 128 sessions (*thick black curve*), the psychometric curves were resampled at 100 uniformly distributed query points ranging from 0 to maximal stimulus amplitude. **B.** To compare neural activity on hits and misses matched in stimulus amplitude and number, we binned the uniform distribution of submaximal stimulus amplitude in each session into 15 equal bins. On bins where there were more hits than misses (e.g. stronger stimulus), we randomly selected hit trials equal to the number of miss trials, and vice versa for bins with more misses than hits. We limited the selection of “matched” trials to trials with reaction time at or longer than 100 ms, reflecting the distribution of observed reaction times (see **C**). The matched trials were subdivided into tertiles based on stimulus amplitude; referred to as *subthreshold, threshold* and *suprathreshold* bins. This accounts for the variation in perceptual threshold between sessions. Unless otherwise noted, hits and misses denote stimulus amplitude matched hits and misses. **C.** *Left* Mean reaction time (RT) on hit trials across all sessions, defined as the time between the sensory onset (also the start of reward window) and the first lick. Cumulative distribution function (CDF) of reaction time on each stimulus amplitude bin shows a sharp reaction time probability inflection at 100 ms, and a median at ~250ms. *Right* The interquartile range (IQR) of RT, for each stimulus amplitude bin. Although the reaction time accelerates slightly with stronger stimulus amplitude, the reaction time variance explained by stimulus amplitude is small relative to the variance within each stimulus amplitude bin. **D.** Extracellular electrophysiology was recorded during the task with chronic drive-able tetrodes from SI ‘barrel’ neocortex. The obtained single units can be classified into fast-spiking units (FS, N=188) and regular-spiking units (putative excitatory neurons; RS, N=278) based on their waveforms (*left*). Specifically, single units where the time between peak and trough ≤ 0.4 ms were classified as FS, and > 0.4 ms as RS (*right*). **E.** Sensory FS (sFS, *gray;* N=69) were defined as those that had a significantly higher firing rate 0 – 100 ms poststimulus after sensory onset on maximal stimulus amplitude trials compared to zero stimulus amplitude (catch) trials (ideal observer analysis, significance determined by 95% bootstrapped confidence interval, CI). In this ideal observer analysis, the area under the receiver operating characteristic (ROC) curve (AUC) is referred to as the stimulus probability (SP). N=5 FS showed significant suppression (*black*). FS that failed to show a significant difference were classified as ‘non-sensory’ (nsFS N=114). **F.** Peristimulus time histograms (PSTH) on matched hit (*blue*) compared to matched miss trials (*red*).PSTHs were calculated with 1 ms bin resolution, then convolved with a Gaussian filter (standard deviation 3 ms, filter length 21 ms). For each FS, the PSTH was averaged across trials and multiplied by 1000, to convert spike rates to Hz. The mean across FS in each subpopulation is shown (sFS *top*,γnsFS *middle* and plnsFS *bottom). Inset* shows firing rate prestimulus (−1000 – 0 ms), poststimulus early (0 – 50 ms) and late (50 – 100 ms) periods, on hits and misses (Mean ± SEM, asterisk if p<0.05 for Wilcoxon signed rank significance tests). **G.** Neurometric curves plotting stimulus evoked rate in the first 100 ms relative to reward/catch window onset, across stimulus amplitude bins on hits/false alarms (*blue)* and misses/correct rejections (*red*). Note that the first 100 ms is always before reaction time, as we only selected hit trials that had reaction time ≥100ms. Error bars indicate standard error of the mean (SEM). Across the entire stimulus range, the sFS response was significantly higher on hit relative to miss trials (Wilcoxon signed rank test, Bonferroni-Holm correction for multiple comparison, * p<0.05). In addition, sFS neurometric curve monotonically increased as the stimulus amplitude increased.

**Supplemental Figure 2.**
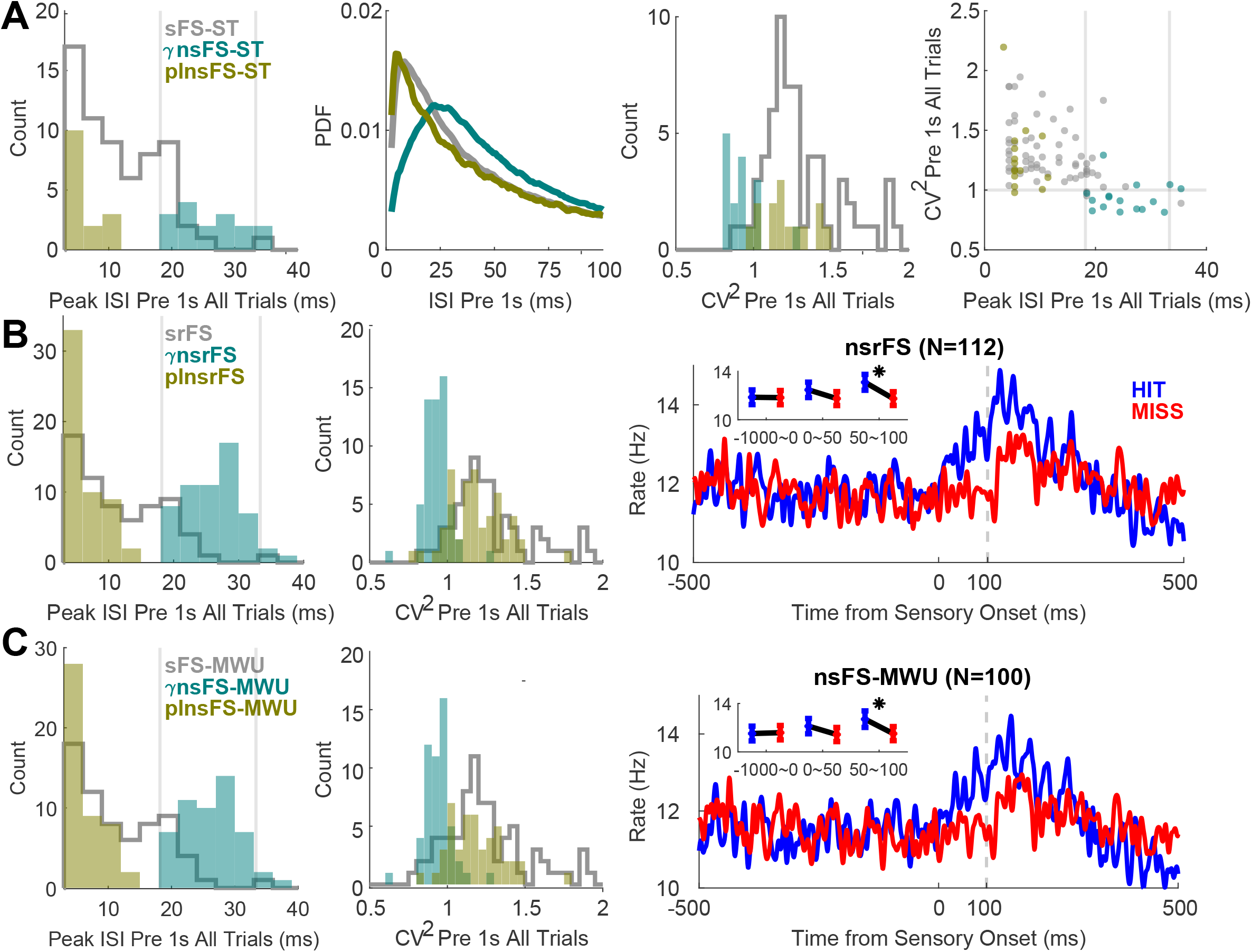
**A.** Same as **Figure 1A-D**, but limited to FS from tetrodes with sensory responsive MUA (ideal observer analysis, significance determined by 95% bootstrapped CI). The restriction to sensory tetrodes (ST) yielded N=64 sFS, 16 γnsFS, and 15 plnsFS. **B-C.** Two alternative definitions of sensory responsiveness are explored. Results are robust in these alternative definitions. Peak ISI histogram (*left*, as in **Figure 1A**), CV^2^ histogram (*center*, as in **Figure 1C**), and PSTH of non-sensory responsive FS for each definition (*right*, as in **Supplemental Figure 1F**) are plotted. **B.** Sensory responsiveness was defined in this analysis based on significant differences between 100 ms prestimulus (−100 – 0 ms) and poststimulus (0 – 100 ms) firing rate (Wilcoxon signed rank test across maximal stimulus amplitude trials, significance at p<0.05). Under this definition, sensory responsive FS showed significantly higher firing rate poststimulus than prestimulus (srFS, N=67; 62 of which overlaped with the sFS definition). Non-sensory responsive FS were not significantly different (nsrFS, N=112; 105 of which overlap with nsFS). **C.** As with the SP based sFS and nsFS definition used throughout, we compared the 0 – 100 ms period on catch trials to maximal stimulus amplitude trials. In this case, instead of bootstrapping ideal observer analysis, sensory responsiveness was determined via a left-tailed Mann-Whitney U-test. sFS-MWU (N=80) are FS with p<0.05, and included all previously defined sFS. nsFS-MWU are not significant in either the left-tailed nor the right-tailed Mann-Whitney U-test (N=100; all of which are nsFS).

**Supplemental Figure 3.**
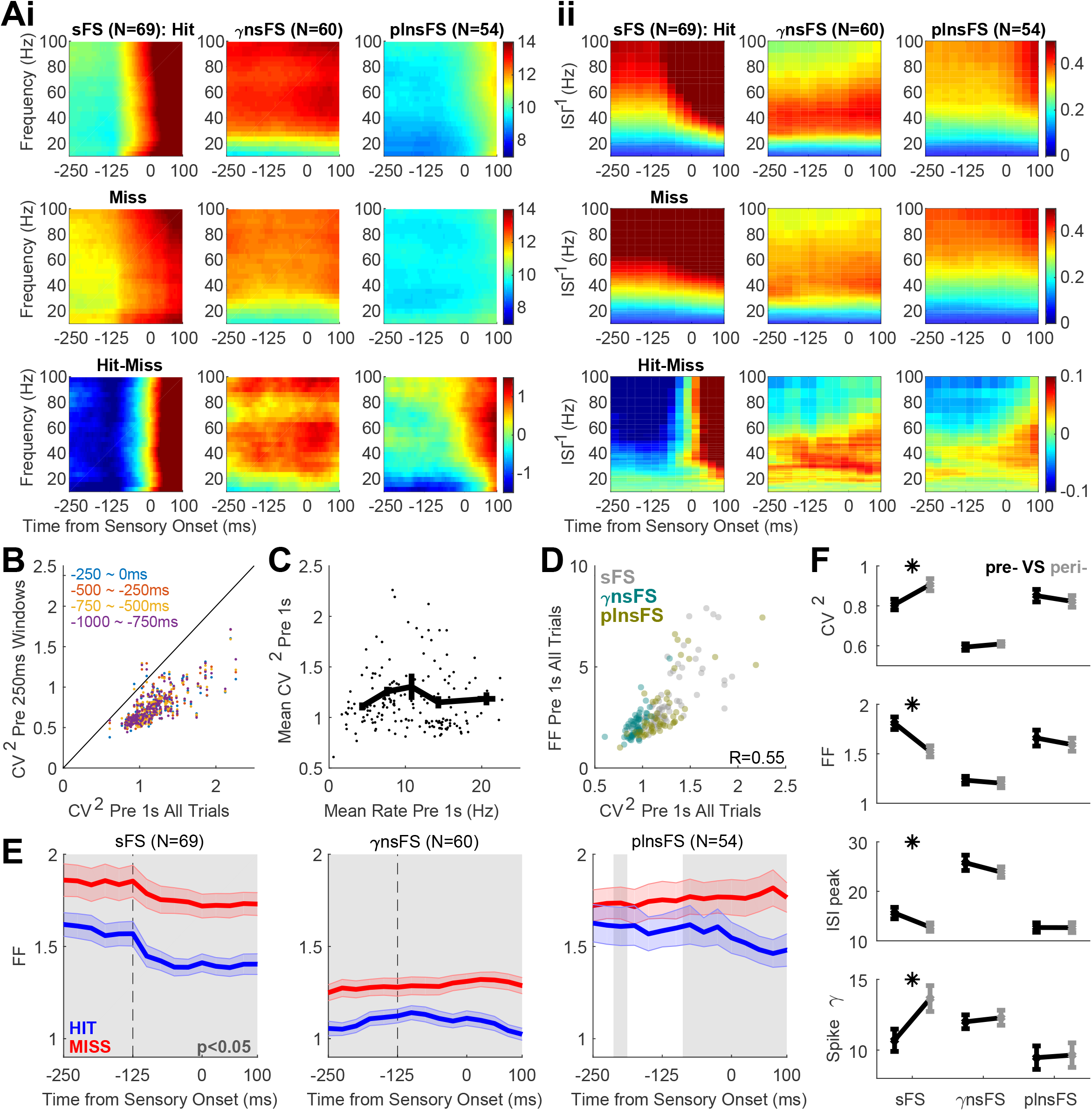
**A.** Side-by-side comparison of spike spectrogram (i) and inverse ISI distribution over time (ii) demonstrate similarity between these measures. (**i**) Spike spectrogram on hits (*top*), misses (*middle*) and hit minus miss (*bottom*); averaged across sFS (*left*),γnsFS (*center*), and plnsFS (*right*). Multitapered spectral power was calculated in 250 ms windows, sliding in 5 ms steps (see *Methods*). (**ii**) Inverse ISI distribution over time on hits (top), misses (*middle*) and hit minus miss (*bottom*); averaged across sFS (*left*),γnsFS (*center*), and plnsFS (*right*). Normalized ISI distribution over time, as described in **Figure 2A**, is represented with inverse ISI on Y-axis. **B.** The numeric values of CV^2^ decrease with smaller time windows, explaining the discrepancy between the CV^2^ values shown in **Figure 1C, D, G** and **Figure 2B**. However, the relative rank order among FS do not change, resulting in high FS-by-FS Spearman’s rank correlation coefficients: Between CV^2^ in the −1000 – 0 ms period (X-axis) and −250 – 0 ms period (Y-axis, *blue*) R_Spearman_=0.78; −500 – −250 ms period (Y-axis, *red*) R_Spearman_=0.77; −750 – −500 ms period (Y-axis, *yellow*) R_Spearman_=0.78; −1000 – −750 ms period (Y-axis, *purple*) R_Spearman_=0.78; and the average of the four non-overlapping 250 ms long windows in the 1 s prestimulus period R_Spearman_=07.9. **C.** CV^2^ does not depend on baseline firing rate. Each point represents an FS. For each FS, the mean firing rate and CV^2^ was calculated across all trials in the 1 s prestimulus. In addition, FS was divided into quintiles based on baseline firing rate, and the mean ± SEM of baseline firing rate (X-axis) and CV^2^ (Y-axis) is plotted in thick black lines. **D.** Fano factor (FF) and CV^2^ are positively correlated across FS, regardless of trial type: R=0.55 on all trials (*left*), hit trials (*center*), and miss trials (*right*). FF is defined as the variance over mean of the firing rate across trials, and was calculated for the 1 s prestimulus period in this figure. **E.** FF calculated in 250 ms windows, shifted in 25 ms steps. Same conventions as in **Figure 2B**: time bin center on X-axis; mean ± SEM across FS in each subgroup on hits (*blue*) and misses (*red*); *gray* shaded regions for significant difference between hits and misses (pointwise Wilcoxon signed rank test p<0.05). sFS (*left*), γnsFS (*center*), and plnsFS (*right*). **F.** Pre- (−400 – −150 ms, *black)* versus peristimulus (−150 – 100 ms, *gray*) comparison of (from *left* to *right*):CV^2^, FF, ISI peak, and spectral power of spike trains in the gamma-band (30 – 55 Hz). Mean ± SEM across sFS (*left* in each plot), γnsFS (*center*), and plnsFS (*right*). Asterisk if p<0.05 for Wilcoxon signed rank test. All four measures of neural dynamics changed significantly for sFS, while none changed for γnsFS nor plnsFS.

**Supplemental Figure 4.**
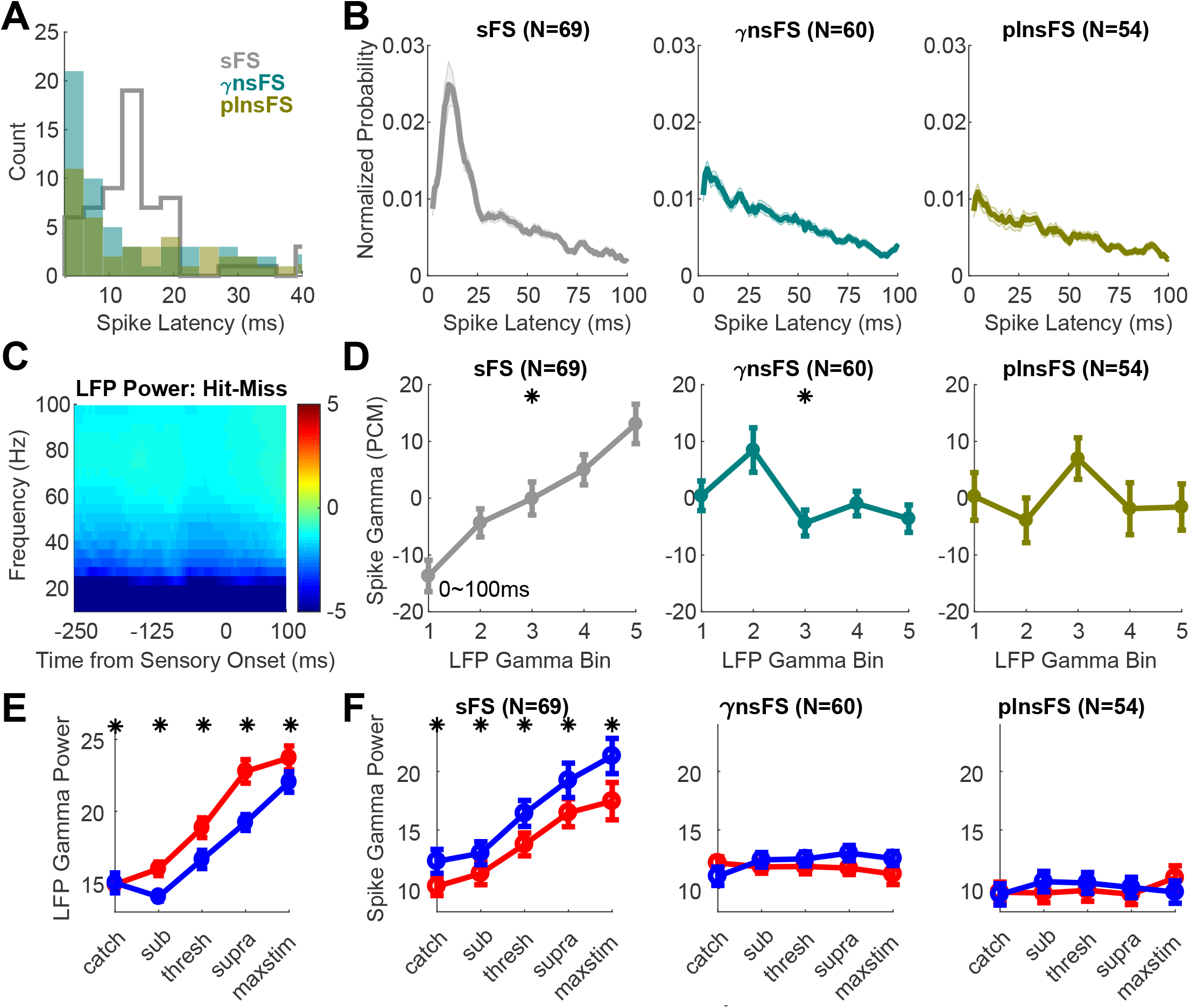
**A.** Normalized histogram of latency to first spike after sensory onset, on maximal stimulus amplitude trials. Histogram counts were calculated in 5 ms bins, 1 ms sliding steps, then normalized by first dividing by the number of steps per bin (i.e. 5) and then by the trial count. Mean ± SEM across sFS (*left*),γnsFS (*center*), and plnsFS (*right)* are depicted. **B.** Peak spike latency histogram (3 ms bins) for sFS (*gray*),γnsFS (*olive*), and plnsFS (*teal*). For each FS, peak spike latency was defined as the first peak in the normalized histogram of spike latency, described in **A**. The sFS distribution showed a distinct peak at 12 – 15 ms bin, whereas both nsFS subtypes showed a monotonic falloff after 3 ms. **C.** The hit minus miss LFP spectrogram, averaged across N=1650 tetrodes. Multitapered spectral power was calculated in 250 ms windows, sliding in 5 ms steps (see *Methods*). **D.** As in **Figure 4D**, but for 0 – 100 ms poststimulus period. **E.** As in **Figure 4B**, but for the LFP gamma power in the in the 0 – 100 ms poststimulus period. **F.** As in **E**, but for the spike gamma power in the 0 – 100 ms poststimulus period. sFS (*left*),γnsFS (*center*),and plnsFS (*right*)

